# Sensory system-specific associations between brain structure and balance

**DOI:** 10.1101/2022.01.17.476654

**Authors:** KE Hupfeld, HR McGregor, CJ Hass, O Pasternak, RD Seidler

## Abstract

Nearly 75% of older adults in the United States report balance problems. Balance difficulties are more pronounced during sensory feedback perturbation (e.g., standing with the eyes closed or on foam). Although it is known that aging results in widespread brain atrophy, less is known about how brain structure relates to balance performance under varied sensory conditions in older age. We measured postural sway of 36 young (18-34 years) and 22 older (66-84 years) adults during four conditions: eyes open, eyes closed, eyes open on foam, and eyes closed on foam. We calculated three summary measures indicating visual, proprioceptive, and vestibular contributions to balance. We also collected *T* _1_-weighted and diffusion-weighted anatomical MRI scans. We aimed to: 1) test for age group differences in brain structure-balance relationships across a range of structural brain measures (i.e., volumetric, surface, and white matter microstructure); and 2) assess how brain structure measures relate to balance, regardless of age. Across both age groups, thinner cortex in multisensory integration regions was associated with greater reliance on visual inputs for balance. Greater gyrification within sensorimotor and parietal cortices was associated with greater reliance on proprioceptive inputs for balance. Poorer vestibular function was correlated with thinner vestibular cortex, greater gyrification within sensorimotor, parietal, and frontal cortices, and lower free water-corrected axial diffusivity in the superior-posterior corona radiata and across the corpus callosum. These results contribute to our scientific understanding of how individual differences in brain structure relate to balance. This has implications for developing brain stimulation interventions to improve balance.

**Significance Statement:** Older age is associated with greater postural sway, particularly when sensory information is perturbed (e.g., by closing one’s eyes). Our work contributes to the field by identifying how individual differences in regional brain structure relate to balance under varying sensory conditions in young and older adults. Across both age groups, lower cortical thickness in sensory integration and vestibular regions, greater gyrification within sensorimotor, parietal, and temporal regions, and lower free water-corrected axial diffusivity in the corpus callosum and corona radiata were related to individual differences in balance scores. We identified brain structures that are associated with specific sensory balance scores; therefore, these results have implications for which brain regions to target in future interventions for different populations.

## Introduction

Balance control declines with older age (e.g., Abrahamova and Hlavač ka 2008; Choy et al. 2003; Colledge et al. 1994; Røgind et al. 2003), and nearly 75% of individuals over the age of 70 in the United States report balance problems (Dillon, 2010). While there are age-related declines to both the peripheral musculoskeletal system (Boelens et al., 2013) and spinal reflexes (Baudry and Duchateau, 2012), degradation of brain structure and function with aging (Seidler et al., 2010) likely also contributes to age-related balance declines. Indeed, studies measuring brain function during standing balance using electroencephalography (Hü lsdü nker et al., 2015; Varghese et al., 2015) and transcranial magnetic stimulation (TMS; Ackermann et al., 1991; Nakazawa et al., 2003) support cortical contributions to balance control (for review, see: Jacobs and Horak 2007; Papegaaij et al. 2014a; Taube et al. 2008).

Postural control is affected by the availability of visual, proprioceptive, and vestibular inputs, which are integrated to signal the body’s orientation and configuration in space (Horak, 2006; Leibowitz and Shupert, 1985; Mahboobin et al., 2005; Peterka, 2002; Shumway-Cook and Horak, 1986). Each of these sensory systems is subject to age-related declines (e.g., reduced receptor numbers; Maki et al. 1999; Patel et al. 2009), and aging also disrupts the relative weighting and integration of their inputs (Colledge et al., 1994; Stelmach et al., 1989; Teasdale et al., 1991; Woollacott et al., 1986). Compared to young adults, older adults experience relatively greater difficulty maintaining their balance during sensory feedback perturbations (e.g., standing with the eyes closed or on foam; Alhanti et al. 1997; Choy et al. 2003; Judge et al. 1995). Here we examined balance across four conditions with varied sensory inputs (i.e., eyes open (EO), eyes closed (EC), eyes open-foam (EOF), and eyes closed-foam (ECF)). This allowed us to characterize individual differences in reliance on visual, proprioceptive, and vestibular inputs.

There is some evidence that brain neurochemistry and function influence balance in older age. For instance, positron emission tomography (PET) measures of striatal dopaminergic denervation (Cham et al., 2007), genetic markers related to dopaminergic transmission (Hupfeld et al., 2018), magnetic resonance spectroscopy metrics of brain antioxidant (glutathione) levels (Hupfeld et al., 2021c), and TMS measures of *γ*-aminobutyric acid (GABA) (Papegaaij et al., 2014b) all correlate with balance performance in older adults. Moreover, functional near-infrared spectroscopy (fNIRS) studies have revealed increased prefrontal brain activity for older adults during standing versus sitting (Mahoney et al., 2016), and increased occipital, frontal, and vestibular cortical activity in older adults during increasingly difficult balance conditions (Lin et al., 2017). These studies provide important insight into the neurochemical and functional correlates of balance control in aging. However, it is widely held that age differences in brain function are at least partially driven by structural brain atrophy (Papegaaij et al., 2014a). Thus, it is important to also understand how individual and age differences in brain structure relate to balance.

Studies of brain structure have shown that poorer balance balance performance in older adults has been linked to larger ventricles (Sullivan et al., 2009; Tell et al., 1998), greater white matter hyperintensity burden (Starr et al., 2003; Sullivan et al., 2009), reduced white matter fractional anisotropy in the corpus callosum (Sullivan et al., 2010; Van Impe et al., 2012), and reduced gray matter volume in the basal ganglia, superior parietal cortex, and cerebellum (Rosano et al., 2007). Other studies have reported no such associations between brain structure and balance in older adults (Ryberg et al., 2007) or opposite relationships between poorer brain structure (e.g., lower basal ganglia gray matter volume) and *better* balance (Boisgontier et al., 2017). Most previous studies investigating associations between brain structure and balance have used only one MRI modality or have focused solely on pathological markers (e.g., white matter hyperintensities instead of ‘normal-appearing’ white matter; Starr et al., 2003; Sullivan et al., 2009). Thus, while this prior work suggests a link between maintenance of brain structure—particularly in sensorimotor regions—in aging and maintenance of postural control, further studies are needed. Moreover, only limited prior work has examined brain structure relationships with sensory-specific balance metrics (Van Impe et al., 2012), though identifying such relationships has implications for better understanding the neural correlates of age-related conditions such as peripheral neuropathy and vestibular dysfunction.

We previously reported on age group differences in brain structure in this cohort (Hupfeld et al., 2021a). In the current study, we addressed two aims: First, we tested for age group differences in the relationship between brain structure and sensory-specific measures of standing balance. As described above, since fNIRS studies show greater prefrontal brain activity for older adults during balance versus sitting (Mahoney et al., 2016), we predicted that greater prefrontal atrophy would correlate more strongly with worse balance scores for the older adults. Second, we determined how sensory-specific measures of standing balance related to brain structure across the whole sample, regardless of age. We hypothesized that, across both young and older adults, we would see functionally specific brain structure-behavior associations in which brain structure in the primary visual, somatosensory, and vestibular cortices would be associated with visual and proprioceptive reliance scores and vestibular function scores, respectively.

## Methods

The University of Florida’s Institutional Review Board provided approval for all procedures performed in this study, and all individuals provided their written informed consent to participate.

### Participants

37 young and 25 older adults participated in this study. Due to the COVID-19 global pandemic, data collection was terminated before we attained the planned sample size for older adults. One young and one older adult were excluded from all analyses because their balance data contained extreme outlier values (*>*5 standard deviations from the group mean). Two older adults were excluded from analyses of the *T* _1_-weighted images: one participant’s head did not fit within the 64-channel coil, so a 20-channel coil was used instead, and we excluded their data due to poor image quality. The other was excluded due to an incidental finding. *T* _1_-weighted images from *n* = 36 young and *n* = 22 older adults were included in analyses. Due to time constraints, diffusion MRI data were not collected for one additional young and two additional older adults; thus, diffusion MRI analyses included data from *n* = 35 young and *n* = 20 older adults.

We screened participants for MRI eligibility and, as part of the larger study, TMS eligibility. We excluded those with MRI or TMS contraindications (e.g., implanted metal, claustrophobia, or pregnancy), history of a neurologic (e.g., stroke, Parkinson’s disease, seizures, or a concussion in the last six months) or psychiatric condition (e.g., active depression or bipolar disorder) or treatment for alcoholism; self-reported smokers; and those who self-reported consuming more than two alcoholic drinks per day on average. Participants were right-handed and were able to stand for at least 30 seconds with their eyes closed. We screened participants for cognitive impairment over the phone using the Telephone Interview for Cognitive Status (TICS-M; de Jager et al., 2003) and excluded those who scored less than 21 of 39 points (which is equivalent to scoring less than 25 points on the Mini-Mental State Exam and indicates probable cognitive impairment (de Jager et al., 2003)). During the first session, we re-screened participants for cognitive impairment using the Montreal Cognitive Assessment (MoCA; Nasreddine et al. 2005). We planned to exclude individuals if they scored less than 23 of 30 points (Carson et al., 2018), but none were excluded for this reason.

### Testing Sessions

We first collected information on demographics (e.g., age, sex, and years of education), self-reported medical history, handedness, footedness, exercise, and sleep. We also collected anthropometric information (e.g., height, weight, and leg length). Participants completed balance testing, followed by an MRI scan approximately one week later. For 24 hours prior to each session, participants were asked not to consume alcohol, nicotine, or any drugs other than the medications they disclosed to us. At the start of each session, participants completed the Stanford Sleepiness Questionnaire, which asks about the number of hours slept the previous night and a current sleepiness rating (Hoddes et al., 1972).

### Balance Testing

Participants completed four balance conditions while instrumented with six Opal inertial measurement units (IMUs; v2; ADPM Wearable Technologies, Inc., Portland, OR, USA). IMUs were placed on the feet, wrists, around the waist at the level of the lumbar spine, and across the torso at the level of the sternal angle. Only the lumbar IMU was used to measure postural sway during standing balance. Participants completed the four-part Modified Clinical Test of Sensory Interaction in Balance (mCTSIB). The mCTSIB has established validity in young and older adults (Alhanti et al., 1997; Cohen et al., 1993; Teasdale et al., 1991) and high retest and inter-tester reliability (Dawson et al., 2018). Participants faced a blank white wall and were instructed to stand as still as possible and refrain from talking for four 30-second trials:

1. eyes open (EO): unperturbed visual, proprioceptive, and vestibular inputs
2. eyes closed (EC): visual input is removed, while proprioceptive and vestibular inputs remain unperturbed
3. eyes open - foam surface (EOF): the foam surface manipulates proprioceptive inputs, but visual and vestibular inputs remain unperturbed
4. eyes closed - foam surface (ECF): visual and proprioceptive cues are compromised, and only vestibular cues are unperturbed

We recorded inertial data during the four trials using MobilityLab software. MobilityLab calculated 25 spatiotemporal features of postural sway using the iSway algorithm (Mancini et al., 2012). We then calculated three summary scores using the 95% ellipse sway area (m^2^/s^4^) variable (i.e., the area of an ellipse covering 95% of the sway trajectory in the coronal and sagittal planes) from each of the four conditions (Fig. 1). Greater postural sway is interpreted as ”worse” standing balance performance (Dewey et al., 2020), as greater postural sway is typically higher for older compared with young adults (e.g., Abrahamova and Hlavač ka, 2008; Colledge et al., 1994; Røgind et al., 2003) and is linked to higher risk of falls (e.g., Laughton et al., 2003; Maki et al., 1994).

**Figure 1:**
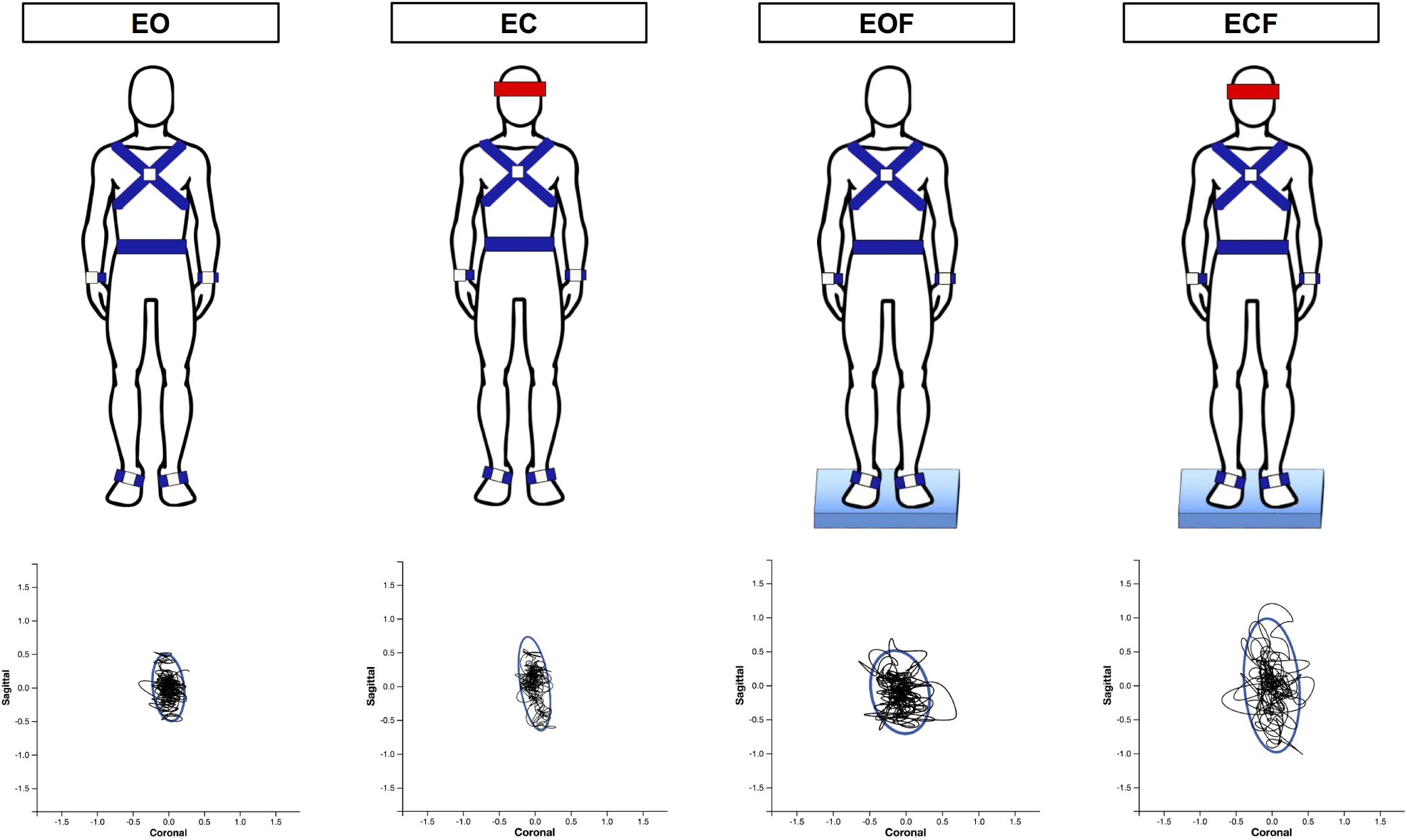
mCTSIB balance conditions. Participants completed four 30-second trials: eyes open (EO), eyes closed (EC), eyes open-foam (EOF), and eyes closed-foam (ECF). Postural sway from each condition was used to calculate the three balance outcome variables, i.e., the visual reliance, proprioceptive reliance, and vestibular function scores. Bottom. Here we depict the sway path (black line) and area (blue oval) for each condition for an exemplar young adult participant. This individual showed greater postural sway as the conditions progressed.

The visual reliance score represents the percent change in postural sway between the eyes closed and the eyes open conditions (considering the foam and firm surface conditions independently and taking the minimum score of the two). Higher scores indicate more difficulty standing still in the absence of visual input. A higher visual reliance score is the result of poorer performance (i.e., more postural sway) during the eyes closed conditions and/or better performance on the eyes open conditions (i.e., less postural sway). Thus, a higher score suggests that the individual is more “reliant” on visual input for balance.

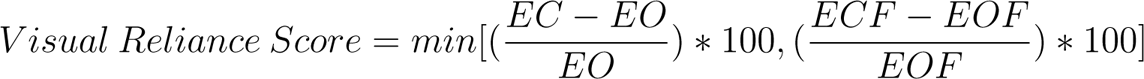

The proprioceptive reliance score represents the percent change between the foam and the firm surface conditions (considering the eyes open and eyes closed conditions independently and taking the minimum of score of the two). Higher scores indicate more difficulty standing still with compromised proprioceptive input. A higher proprioceptive reliance score is the result of poorer performance (i.e., more postural sway) on the foam conditions and/or better performance on the firm surface conditions (i.e., less postural sway). Thus, a higher score suggests that the individual is more “reliant” on proprioceptive input for balance.

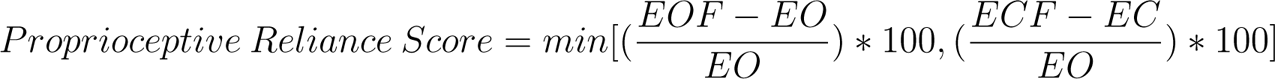

The vestibular function score represents the percent change between the ECF and EO conditions. Higher scores indicate more difficulty standing still when only vestibular input is appropriate and visual / proprioceptive inputs are compromised. Contrary to the scores described above (which represent reliance on visual and proprioceptive inputs, respectively), higher scores here indicate *poorer* vestibular function (Dewey et al., 2020).

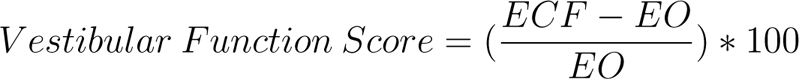

These formulas represent those recommended by APDM (the IMU company) for calculating mCTSIB summary scores (for further details, see: https://support.apdm.com/hc/en-us/articles/217035886-How-are-the-ICTSIB-composite-scores-computed-). For simplicity and to keep with prior literature (Goble et al., 2019, 2020), we will use the interpretation of higher visual and proprioceptive scores indicating more ”reliance” on these two sensory systems for balance. However, it is worth noting that this interpretation might be oversimplified. These scores may also index sensory reweighting and integration more so than reliance on a single sensory modality (Kalron, 2017). We expand on this in the Discussion.

### MRI Scan

We used a Siemens MAGNETOM Prisma 3 T scanner (Siemens Healthcare, Erlangen, Germany) with a 64-channel head coil to collect *T* _1_-weighted and diffusion-weighted scans for each participant. We collected the 3D *T* _1_-weighted anatomical image using a magnetization-prepared rapid gradient-echo (MPRAGE) sequence. The parameters were: repetition time (TR) = 2000 ms, echo time (TE) = 3.06 ms, flip angle = 8°, field of view = 256 × 256 mm^2^, slice thickness = 0.8 mm, 208 slices, voxel size = 0.8 mm^3^. Next, we collected the diffusion-weighted spin-echo prepared echo-planar imaging sequence with the following parameters: 5 *b*_0_ volumes (without diffusion weighting) and 64 gradient directions with diffusion weighting 1000 s/mm^2^, TR = 6400 ms, TE = 58 ms, isotropic resolution = 2 x 2 x 2 mm, FOV = 256 x 256 mm^2^, 69 slices, phase encoding direction = Anterior to Posterior. Immediately prior to this acquisition, we collected 5 *b*_0_ volumes (without diffusion weighting) in the opposite phase encoding direction (Posterior to Anterior) for later use in distortion correction.

### T***_1_***-Weighted Image Processing for Voxelwise Analyses

We used the same *T* _1_-weighted processing steps as described in our previous work (Hupfeld et al., 2021a).

### Gray matter volume

We processed the *T* _1_-weighted scans using the Computational Anatomy Toolbox (CAT12; version r1725; Gaser et al., 2016; Gaser and Kurth, 2017) in MATLAB R2019b. We implemented default CAT12 preprocessing steps. This included segmentation into gray matter, white matter, and cerebrospinal fluid, followed by spatial normalization to standard space using high-dimensional Dartel registration and modulation. Modulation involves multiplying the normalized gray matter segment by its corresponding Jacobian determinant to produce modulated gray matter volume images in standard space. The Jacobian determinant encodes local shrinkage and expansion between subject space and the target image (i.e., standard space template). To increase signal-to-noise ratio, we smoothed the modulated, normalized gray mattersegments using Statistical Parametric Mapping 12 (SPM12, v7771; Ashburner et al., 2014) with an 8 mm full width at half maximum kernel. We entered the smoothed, modulated, normalized gray matter volume maps into the group-level voxelwise statistical models described below. We used CAT12 to calculate total intracranial volume for each participant for later use as a covariate in our group-level statistical analyses.

### Cortical surface metrics

The CAT12 pipeline also extracts surface-based voxelwise morphometry metrics (Dahnke et al., 2013; Yotter et al., 2011a) using a projection-based thickness algorithm that handles partial volume information, sulcal blurring, and sulcal asymmetries without explicit sulcus reconstruction (Dahnke et al., 2013; Yotter et al., 2011a). We extracted four surface metrics: 1) cortical thickness: the width of the cortical gray matter between the outer surface (i.e., the gray matter-cerebrospinal fluid boundary) and the inner surface (i.e., the gray matter-white matter boundary) (Dahnke et al., 2013); 2) cortical complexity: fractal dimension, a metric of folding complexity of the cortex (Yotter et al., 2011b); 3) sulcal depth: the Euclidean distance between the central surface and its convex hull (Yun et al., 2013); and 4) gyrification index: a metric based on the absolute mean curvature, which quantifies the amount of cortex buried within the sulcal folds as opposed to the amount of cortex on the “outer” visible surface (Luders et al., 2006). We resampled and smoothed the surfaces at 15 mm for cortical thickness and 20 mm for the three other metrics. We entered these resampled and smoothed surface files into our group-level voxelwise statistical models.

### Cerebellar volume

Similar to our past work (Hupfeld et al., 2021b; Salazar et al., 2020, 2021), we applied specialized preprocessing steps to the cerebellum to produce cerebellar volume maps, with improved normalization of the cerebellum (Diedrichsen, 2006; Diedrichsen et al., 2009). We entered each participant’s whole-brain *T* _1_-weighted image into the CEREbellum Segmentation (CERES) pipeline (Romero et al., 2017). We used a binary mask from each participant’s CERES cerebellar segmentation to extract their cerebellum from their whole-brain *T* _1_-weighted image. We used rigid, affine, and Symmetric Normalization (SyN) transformation procedures in the Advanced Normalization Tools package (ANTs; v1.9.17; Avants et al., 2010, 2011) to warp (in a single step) each participant’s extracted subject space cerebellum to a 1 mm cerebellar template in standard space, the Spatially Unbiased Infratentorial Template (SUIT) (Diedrichsen, 2006; Diedrichsen et al., 2009). The flowfields used to warp native cerebellar segments directly to SUIT space were additionally used to calculate the Jacobian determinant image, using ANTs’ *CreateJacobianDeterminantImage.sh* function. We multiplied each normalized cerebellar segment by its corresponding Jacobian determinant to produce modulated cerebellar images in standard space. To increase signal-to-noise ratio, we smoothed the modulated, normalized cerebellar images using a kernel of 2 mm full width at half maximum and entered the resulting cerebellar volume maps into our group-level voxelwise statistical models.

### Diffusion-Weighted Image Processing for Voxelwise Analyses

We used the same diffusion-weighted processing steps as described in detail in our previous work (Hupfeld et al., 2021a).

### Diffusion preprocessing

We corrected images for signal drift (Vos et al., 2017) using the ExploreDTI graphical toolbox (v4.8.6; www.exploredti.com; Leemans et al., 2009) in MATLAB (R2019b). Next, we used the FMRIB Software Library (FSL; v6.0.1; Jenkinson et al., 2012; Smith et al., 2004) processing tool, *topup*, to estimate the susceptibility-induced off-resonance field (Andersson et al., 2003). This yielded a single corrected field map for use in eddy current correction. We used FSL’s *eddy cuda* to simultaneously correct the data for eddy current-induced distortions and both inter- and intra-volume head movement (Andersson and Sotiropoulos, 2016).

### FW correction and tensor fitting

We implemented a custom free-water (FW) imaging algorithm (Pasternak et al., 2009) in MATLAB. This algorithm estimates FW fractional volume and FW-corrected diffusivities by fitting a two-compartment model at each voxel (Pasternak et al., 2009). The FW compartment reflects the proportion of water molecules with unrestricted diffusion and is quantified by the fractional volume of this compartment. FW fractional volume ranges from 0 to 1; FW = 1 indicates that a voxel is filled with freely diffusing water molecules (e.g., within the ventricles). The tissue compartment models FW-corrected indices of water molecule diffusion within or in the vicinity of white matter tissue, quantified by diffusivity (FAt, RDt, and ADt). These metrics (FW, FAt, RDt, ADt) are provided as separate voxelwise maps.

### Tract-Based Spatial Statistics

We applied FSL’s tract-based spatial statistics (TBSS) processing steps to prepare the data for voxelwise analyses across participants (Smith et al., 2006). TBSS was selected because it avoids problems associated with suboptimal image registration between participants and does not require spatial smoothing. TBSS uses a carefully-tuned nonlinear registration and projection onto an alignment-invariant tract representation (i.e., the mean FA skeleton); this process improves the sensitivity, objectivity, and interpretability of analyses of multi-subject diffusion studies. We used the TBSS pipeline as provided in FSL. This involves eroding the FA images slightly and zeroing the end slices, then bringing each subject’s FA data into standard space using the nonlinear registration tool FNIRT (Andersson et al., 2007b,a). A mean FA image is then calculated and thinned to create a mean FA skeleton. Each participant’s aligned FA data is then projected onto the group mean skeleton. Lastly, we applied the same nonlinear registration to the FW, FAt, RDt, and ADt maps to project these data onto the original mean FA skeleton. Ultimately, these TBSS procedures resulted in skeletonized FW, FAt, ADt, and RDt maps in standard space for each participant. These were the maps that we entered in our group-level voxelwise statistical models.

### Ventricular Volume Calculation

CAT12 automatically calculates the inverse warp, from standard space to subject space, for the Neuromorphometrics (http://Neuromorphometrics.com) volume-based atlas. We isolated the lateral ventricles from this atlas in subject space. We visually inspected the ventricle masks overlaid onto each participant’s *T* _1_-weighted image in ITK-SNAP and hand corrected the ROI mask if needed (Yushkevich et al., 2006). Using *fslstats*, we extracted the number of voxels in each ventricular mask in subject space and calculated the mean image intensity within the ventricles in the subject space cerebrospinal fluid segment. We then calculated each lateral ventricular volume, in mL, as: (number of voxels in the ventricular mask)*(mean intensity of the cerebrospinal fluid probabilistic map within the ROI mask)*(volume/voxel). In subsequent statistical analyses, we used the average of the left and right side structures for each ROI, and we entered these ROI volumes as a percentage of total intracranial volume to account for differences in head size.

### Statistical Analyses

#### Participant characteristics, testing timeline, and balance

We conducted all statistical analyses on the demographic and balance data using R (v4.0.0; R Core Team, 2013). We conducted nonparametric two-sided Wilcoxon rank-sum tests for age group differences in demographics, physical characteristics, and session timeline variables. We used a Pearson chi-square test to check for differences in the sex distribution within each age group. We used three linear models to test for age group differences in the balance scores (i.e., visual, proprioceptive, and vestibular), controlling for sex. We applied the Benjamini-Hochberg false discovery rate (FDR) correction to the *p* values for the age group predictor (Benjamini and Hochberg, 1995).

#### Voxelwise Statistical Models

We tested the same voxelwise models for each of the imaging modalities. In each case, we defined the model using SPM12 and then re-estimated the model using the Threshold-Free Cluster Enhancement toolbox (TFCE; http://dbm.neuro.uni-jena.de/tfce) with 5,000 permutations. This toolbox provides non-parametric estimation using TFCE for models previously estimated using SPM parametric designs. Statistical significance was determined at *p* < 0.05 (two-tailed) and family-wise error (FWE) corrected for multiple comparisons. In each of the below models, we set the brain structure map as the outcome variable. In the gray matter volume models only, we set the absolute masking threshold to 0.1 (Gaser and Kurth, 2017) and used an explicit gray matter mask that excluded the cerebellum (because we analyzed cerebellar volume separately from “whole brain” gray matter volume).

#### Age group differences in brain structure

We previously reported the results of two-sample t-tests for age group differences in brain structure (Hupfeld et al., 2021a).

#### Interaction of age group and balance scores

First, we tested for regions in which the relationship between brain structure and balance performance differed between young and older adults. We ran independent samples t-tests and included the balance scores for young and older adults as covariates of interest. We tested for regions in which the correlation between brain structure and balance performance differed between young and older adults (i.e., for statistical significance in the interaction term). We controlled for sex in all models and also for head size (i.e., total intracranial volume, as calculated by CAT12) in the gray matter and cerebellar volume models.

#### Whole group correlations of brain structure with balance scores

Next, we conducted a linear regression omitting the age group*balance score interaction term, to test for regions of association between brain structure and balance performance, regardless of age or sex. That is, in each of these models, we included the whole cohort and controlled for age and sex (but did not include an age group predictor or interaction term). In the gray matter and cerebellar volume models, we also controlled for head size.

### Ventricular volume statistical models

We carried out linear models in R to test for relationships between ventricular volume and balance, controlling for age and sex. We then ran linear models testing for an interaction between age group and balance scores, controlling for sex. In each case, we FDR-corrected the *p* values for the predictor of interest (i.e., balance score or the interaction term, respectively; Benjamini and Hochberg, 1995).

### Multiple regression to fit the best model of vestibular function scores in older adults

We used a stepwise multivariate linear regression to directly compare the predictive strength of the brain structure correlates of balance scores identified by the analyses described above. We ran one model for the vestibular function scores (as the visual and proprioceptive reliance scores did not produce more than one resulting brain structure measure). We included as predictors age, sex and values from the peak result coordinate for each model that indicated a statistically significant relationship between brain structure and vestibular function scores. We used *stepAIC* (Venables et al., 1999) to produce a final model that retained only the best predictor variables; *stepAIC* selects a maximal model based on the combination of predictors that produces the smallest Akaike information criterion (AIC). This stepwise regression approach allowed us to fit the best model using brain structure to predict vestibular function scores.

## Results

### Age Differences in Participant Characteristics, Testing Timeline, and Balance

There were no significant differences between the age groups for most demographic variables, including sex, handedness, footedness, and alcohol use (Table 1). The older adults had higher body mass indices and exercised less compared to the young adults. The older adults reported a greater fear of falling and less balance confidence. There were no age group differences in the number of days elapsed between the two testing sessions or in the difference in start time for the sessions.

**Table 1:** Participant characteristics and testing timeline

| Variables | Young adult<br>median (IQR) | Older adult<br>median (IQR) | W or $\chi^2$ | FDR<br>corr. $p$ | Effect<br>size<sup>a</sup> |
| --- | --- | --- | --- | --- | --- |
| Demographics |  |  |  |  |  |
| Sample size | 36 | 22 |  |  |  |
| Age (years) | 21.75 (2.36) | 72.58 (9.72) |  |  |  |
| Sex | 19 F; 17 M | 12 F; 10 M | 0.02 | 0.896 |  |
| Physical characteristics and fitness |  |  |  |  |  |
| Handedness laterality score<sup>b</sup> | 85.17 (25.42) | 100.00 (24.55) | 329.50 | 0.423 | -0.14 |
| Footedness laterality score<sup>b</sup> | 100.00 (22.22) | 100.00 (138.39) | 452.50 | 0.455 | -0.13 |
| Body mass index (kg/m<sup>2</sup>) | 22.76 (5.67) | 25.92 (3.76) | 175.00 | **0.006\*\*** | -0.46 |
| Leisure-time physical activity<sup>c</sup> | 46.00 (38.00) | 29.00 (21.00) | 551.00 | **0.017\*** | -0.37 |
| Balance and fear of falling |  |  |  |  |  |
| Balance confidence<sup>d</sup> | 97.81 (3.61) | 94.07 (4.38) | 595.50 | **0.010\*\*** | -0.41 |
| Fear of falling<sup>d</sup> | 17.00 (3.00) | 19.00 (2.00) | 224.50 | **0.017\*** | -0.36 |
| Education and cognition |  |  |  |  |  |
| Years of education | 15.00 (3.00) | 16.00 (4.25) | 226.00 | **0.017\*\*** | -0.35 |
| MoCA score | 28.00 (3.25) | 27.00 (2.75) | 517.50 | 0.114 | -0.25 |
| Alcohol use |  |  |  |  |  |
| AUDIT score<sup>e</sup> | 2.00 (3.50) | 1.00 (3.75) | 495.00 | 0.219 | -0.21 |
| Hours of sleep |  |  |  |  |  |
| Behavioral session | 7.00 (1.62) | 7.50 (1.50) | 343.00 | 0.656 | -0.07 |
| MRI session | 7.00 (1.62) | 7.00 (1.00) | 319.50 | 0.382 | -0.16 |
| Testing timeline<sup>f</sup> |  |  |  |  |  |
| Behavioral vs. MRI session (days) | 3.50 (6.25) | 4.50 (4.75) | 357.00 | 0.656 | -0.08 |
| Behavioral vs. MRI start (hours) | 1.33 (1.41) | 1.23 (1.10) | 419.50 | 0.767 | -0.05 |

*Note:* In the second and third columns, we report the median ± interquartile range (IQR) for each age group in all cases except for sex. For sex, we report the number of males and females in each age group. In the fourth and fifth columns, for all variables except sex, we report the result of a nonparametric two-sample, two-sided Wilcoxon rank-sum test. For sex, we report the result of a Pearson's chi-square test for differences in the sex distribution within each age group. All participants with $T_1$ -weighted scans are included in the comparisons in this table. However, we excluded several individuals from the diffusion-weighted image analyses (see Methods). $P$ values were FDR-corrected (Benjamini and Hochberg, 1995) across all models included in this table. \* $p < 0.05$ , \*\* $p < 0.01$ . Significant $p$ values are bolded.
<sup>a</sup>In the sixth column, we report the nonparametric effect size as described by (Rosenthal et al., 1994; Field et al., 2012).
<sup>b</sup>We calculated handedness and footedness laterality scores using two self-report surveys: the Edinburgh Handedness Inventory (Oldfield, 1971) and the Waterloo Footedness Questionnaire (Elias et al., 1998).
<sup>c</sup>We assessed self-reported physical activity using the Godin Leisure-Time Exercise Questionnaire (Godin et al., 1985).
<sup>d</sup>Participants self-reported Activities-Specific Balance Confidence scores (Powell and Myers, 1995) and fear of falling using the Falls Efficacy Scale (Tinetti et al., 1990).
<sup>e</sup>Participants self-reported alcohol use on the Alcohol Use Disorders Identification Test (AUDIT; Piccinelli, 1998).
<sup>f</sup>Here we report the days between the testing sessions and the hours between the start time of the testing sessions.

No age group differences emerged for visual reliance scores. That is, young and older adults showed a similar increase in postural sway under the eyes closed compared to the eyes open balance conditions (i.e., visual reliance score; Table 2; Fig. 2A). Older adults had higher proprioceptive reliance compared to young adults, exhibiting greater postural sway during the foam versus firm surface conditions (i.e., proprioceptive reliance score; Fig. 2B). Further, older adults had poorer (i.e., higher) vestibular function scores compared to the young adults. That is, older adults exhibited greater postural sway during the ECF versus EO conditions, indicating poorer vestibular function (i.e., poorer performance when visual and proprioceptive inputs were compromised and only vestibular input was available; Fig. 2C).

**Figure 2:**
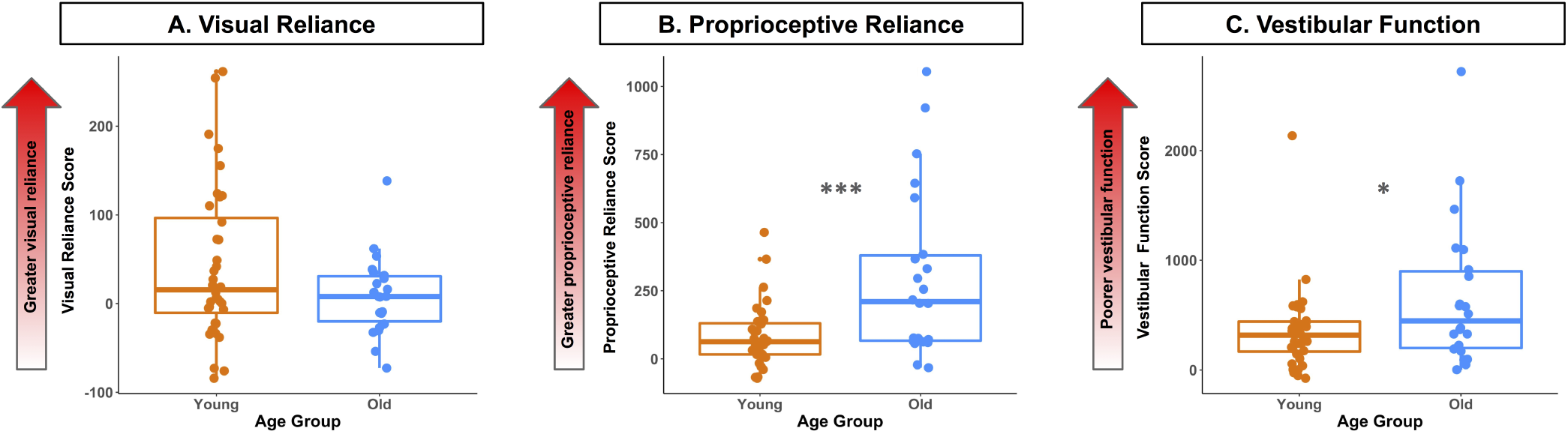
Age group differences in balance composite scores. Balance scores are shown for the older (blue) and younger (orange) adults. The red arrows point in the direction of higher scores. Higher scores indicate a greater reliance on visual (A) and proprioceptive (B) inputs for maintaining standing balance, or poorer vestibular function (C).

**Table 2:** Age differences in balance scores

| Mean (SD) |  | Predictors | Estimates (SE) | t | FDR<br>Corr. *p* | R<sup>2</sup> |
| --- | --- | --- | --- | --- | --- | --- |
| Visual reliance |  | (Intercept) | 42.55 (12.41) | 3.43 |  |  |
| Young: 42.89 (86.99) | Old: 8.71 (44.44) | Age group (Old) | -34.39 (20.13) | -1.71 | 0.093 |  |
|  |  | Sex (Male) | 6.01 (9.79) | 0.61 |  | 0.06 |
| Proprioceptive reliance |  | (Intercept) | 82.64 (35.28) | 2.34 |  |  |
| Young: 82.07 (115.39) | Old: 301.56 (308.63) | Age group (Old) | 219.85 (57.24) | 3.84 | **<0.001\*\*\*** |  |
|  |  | Sex (Male) | -10.21 (27.84) | -0.37 |  | 0.21 |
| Vestibular function |  | (Intercept) | 348.78 (83.92) | 4.16 |  |  |
| Young: 348.75 (373.56) | Old: 654.36 (655.86) | Age group (Old) | 305.64 (136.15) | 2.25 | **0.043\*** |  |
|  |  | Sex (Male) | -0.63 (66.22) | -0.01 |  | 0.08 |

Note: On the left side, we report mean (standard deviation) for the young and older age groups. On the right side, we report the results of three linear models testing for age group differences in each balance score, controlling for sex. *P* values for the age group predictor were FDR-corrected (Benjamini and Hochberg, 1995). SD = standard deviation; SE = standard error. \* $p_{FDR-corr} < 0.05$ , \*\*\* $p_{FDR-corr} < 0.001$ . Significant *p* values are bolded.

### Age Group Differences in Brain Structure

Our recent publication provides a detailed report of age group differences in brain structure in this cohort (Hupfeld et al., 2021a). Overall, we found evidence of widespread cortical and cerebellar atrophy for older compared with young adults across the examined volumetric, surface, white matter microstructure, and ventricle metrics. Interestingly, we identified the most prominent age differences in several metrics (i.e., gray matter volume and cortical thickness) in the sensorimotor cortices, and comparatively less age difference in these metrics in the frontal cortices. Refer to Hupfeld et al. (2021a) for further details.

### No Age Differences in the Relationship of Brain Structure with Balance

Across all brain structure metrics, there were no age differences in the relationship between the balance scores and brain structure. That is, there was no interaction of age group and balance scores; therefore, our second set of statistical analyses did not include an interaction term and instead aimed to identify relationships between brain structure and balance scores across the whole cohort (regardless of age).

### Brain Structure Correlates of Balance Scores

There were no relationships between gray matter volume, cortical complexity, sulcal depth, or cerebellar volume and balance performance across the whole cohort. Thinner cortex (i.e., ”worse” brain structure) within a region encompassing portions of the right cingulate gyrus (isthmus), precuneus, and lingual gyrus was associated with higher visual reliance scores (Fig. 3; Table 3). That is, those individuals who had the thinnest cortex in these regions also showed the greatest increase in postural sway between conditions with the eyes closed compared with open (indicating greater reliance on visual inputs for balance). In addition, thinner cortex within two regions encompassing portions of the left supramarginal and postcentral gyri and the bank of the left superior temporal sulcus was associated with poorer vestibular function scores (Fig. 3; Table 3). That is, those individuals who had the thinnest cortex in these regions also exhibited the most postural sway during the ECF relative to the EO condition (indicating poorer vestibular function).

**Figure 3:**
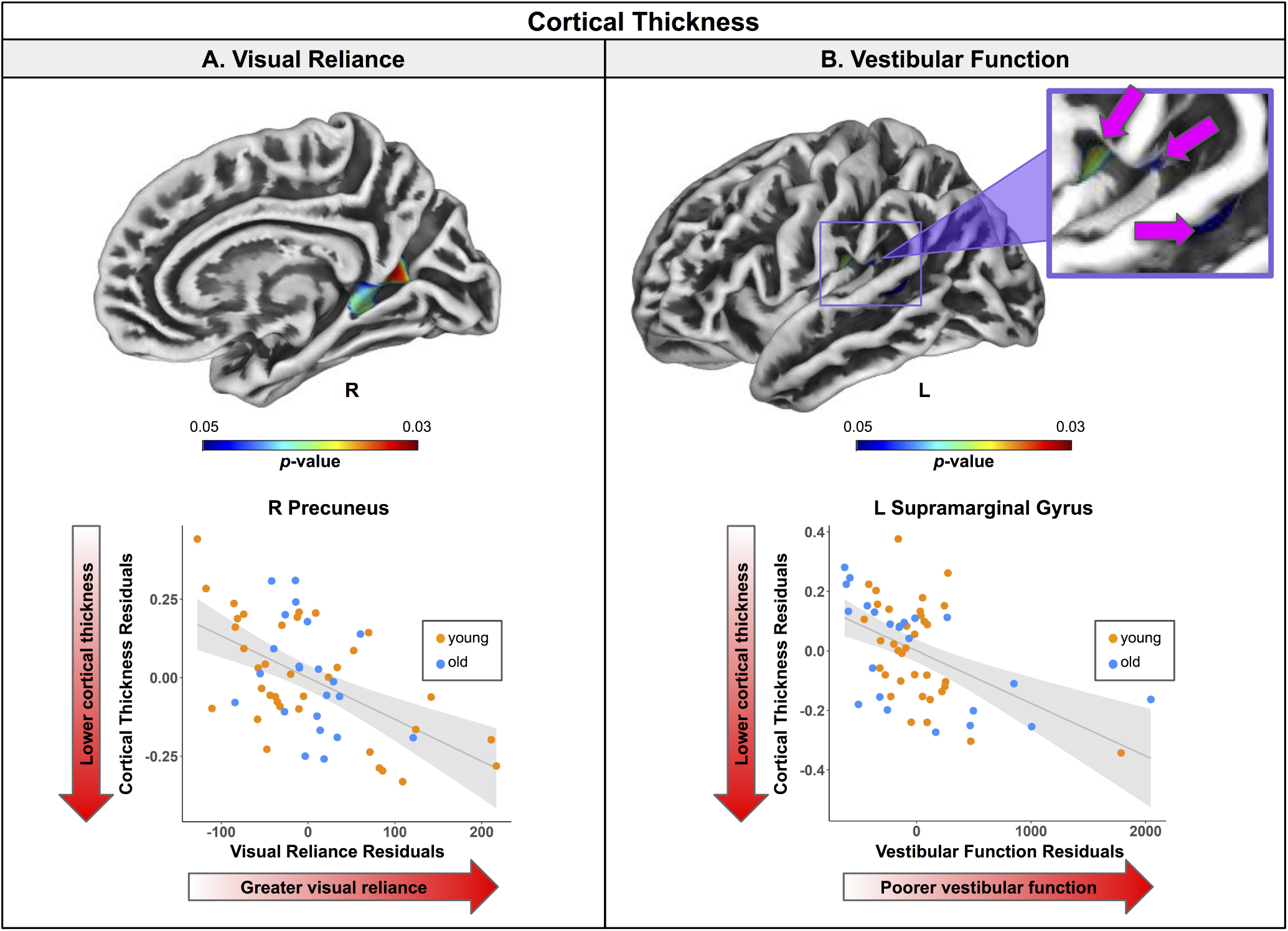
Regions of correlation between cortical thickness and balance scores. Top. Regions showing statistically significant (*p_FWE−corr_* < 0.05) relationships between cortical thickness and vision (left) and vestibular (right) balance scores. Warmer colors indicate regions of stronger correlation. Results are overlaid onto CAT12 standard space templates. L = left hemisphere; R = right hemisphere. Bottom. Surface values for the peak result coordinate for each model are plotted against balance scores to illustrate examples of the relationships identified by the voxelwise statistical tests. The fit line and confidence interval shading are included only to aid visualization of these relationships. We plotted the residuals instead of the raw values here to adjust for the effects of the age and sex covariates included in each model.

**Table 3:** Regions of correlation between cortical thickness and balance scores

| Region | Overlap of Atlas Region | TFCE Level |  |
| --- | --- | --- | --- |
| | | Extent ($k_E$) | $p_{FWE-corr}$ |
| Visual reliance |  |  |  |
| R cingulate gyrus (isthmus) | 43% | 344 | **0.030\*** |
| R precuneus | 39% | — | — |
| R lingual gyrus | 15% | — | — |
| R pericalcarine cortex | 1% | — | — |
| Vestibular function |  |  |  |
| L supramarginal gyrus | 69% | 188 | **0.038\*** |
| L postcentral gyrus | 31% | — | — |
| L superior temporal sulcus (bank) | 100% | 55 | **0.049\*** |

Note: Here we list all atlas regions from the Desikan-Killiany DK40 atlas (Desikan et al., 2006) that overlapped with each resulting cluster. We do not list volumetric (e.g., MNI space) coordinates in this table because volumetric coordinates cannot be mapped directly onto cortical surfaces. L = left; R = right. $*p_{FWE-corr} < 0.05$ . Significant $p$ values are bolded.

Higher gyrification index (i.e., ”better” brain structure; Luders et al., 2006) within two large clusters encompassing portions of the left sensorimotor, parietal, supramarginal, paracentral, and frontal cortices and precuneus was associated with higher proprioceptive reliance scores (Fig. 4; Table 4). That is, those individuals with the highest gyrification index in these regions also showed the greatest increase in postural sway for conditions using the foam compared to the firm surfaces (indicating greater reliance on proprioceptive inputs for balance).

**Figure 4:**
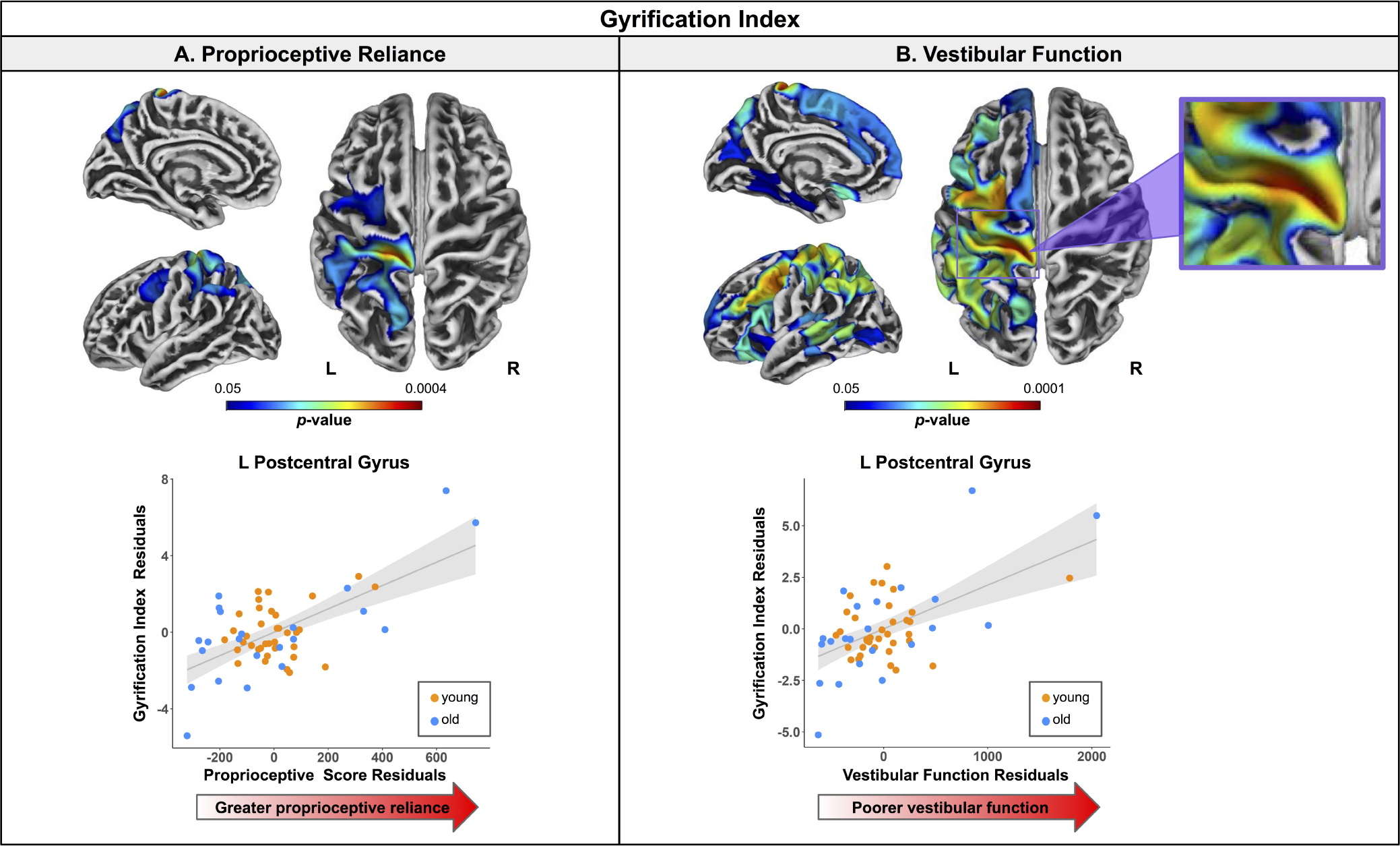
Regions of correlation between gyrification index and balance scores. Top. Regions showing statistically significant (*p_FWE−corr_* < 0.05) relationships between gyrification index and proprioceptive (A) and vestibular (B) balance scores. Warmer colors indicate regions of stronger correlation. Results are overlaid onto CAT12 standard space templates. L = left hemisphere; R = right hemisphere. Bottom. Surface values for the peak result coordinate for each model are plotted against balance score to illustrate examples of the relationships identified by the voxelwise statistical tests. The fit line and confidence interval shading are included only to aid visualization of these relationships. We plotted the residuals instead of the raw values here to adjust for the effects of the age and sex covariates included in each model.

**Table 4:** Regions of correlation between gyrification index and balance scores

| Region | Overlap of Atlas Region | TFCE Level |  |
| --- | --- | --- | --- |
| | | Extent ($k_E$) | $p_{FWE-corr}$ |
| Proprioceptive reliance |  |  |  |
| L postcentral gyrus | 30% | 2555 | **<0.001\*\*\*** |
| L superior parietal cortex | 29% | — | — |
| L supramarginal gyrus | 19% | — | — |
| L precentral gyrus | 14% | — | — |
| L precuneus | 5% | — | — |
| L inferior parietal cortex | 2% | — | — |
| L paracentral gyrus | 1% | — | — |
| L caudal middle frontal gyrus | 54% | 800 | **0.02\*** |
| L precentral gyrus | 32% | — | — |
| L superior frontal gyrus | 14% | — | — |
| Vestibular function |  |  |  |
| L superior frontal gyrus | 12% | 13292 | **<0.001\*\*\*** |
| L superior parietal cortex | 11% | — | — |
| L precentral gyrus | 9% | — | — |
| L postcentral gyrus | 9% | — | — |
| L supramarginal gyrus | 7% | — | — |
| L inferior parietal cortex | 6% | — | — |
| L rostral middle frontal gyrus | 6% | — | — |
| L caudal middle frontal gyrus | 5% | — | — |
| L insula | 4% | — | — |
| L lateral orbitofrontal cortex | 4% | — | — |
| L superior temporal cortex | 4% | — | — |
| L superior temporal sulcus (bank) | 3% | — | — |
| L pars opercularis | 3% | — | — |
| L middle temporal gyrus | 3% | — | — |
| L precuneus | 2% | — | — |
| L pars triangularis | 2% | — | — |
| L cuneus | 2% | — | — |
| L lateral occipital cortex | 2% | — | — |
| L lateral paracentral gyrus | 1% | — | — |
| L caudal anterior cingulate gyrus | 1% | — | — |
| L medial orbitofrontal gyrus | 1% | — | — |
| L lingual gyrus | 32% | 961 | **0.031\*** |
| L fusiform gyrus | 30% | — | — |
| L parahippocampal gyrus | 28% | — | — |
| L entorhinal cortex | 7% | — | — |
| L cingulate gyrus (isthmus) | 3% | — | — |
| R lateral orbitofrontal cortex | 100% | 38 | **0.049\*** |

Note: Here we list all atlas regions from the Desikan-Killiany DK40 atlas (Desikan et al., 2006) that overlapped with each resulting cluster. We do not list volumetric (e.g., MNI space) coordinates in this table because volumetric coordinates cannot be mapped directly onto cortical surfaces. L = left; R = right. \* $p_{FWE-corr} < 0.05$ ; \*\*\* $p_{FWE-corr} < 0.001$ . Significant $p$ values are bolded.

In addition, higher gyrification index within a large region spanning portions of the frontal, temporal, and parietal cortices was associated with poorer vestibular function scores (Fig. 4; Table 4). That is, those individuals who had the highest gyrification index in these regions also exhibited the most postural sway during the ECF relative to the EO condition (indicating poorer vestibular function). This relationship between ”better” brain structure and worse vestibular function is seemingly contradictory, though these resulting regions did *not* include the so-called vestibular cortices (Lopez et al., 2012; zu Eulenburg et al., 2012). It could be that those with poorer vestibular function rely more on other brain regions for balance, as compensation. We expand on this idea in the Discussion.

Poorer vestibular function scores were also associated with lower ADt (i.e., typically interpreted as ”worse” brain structure; Bennett et al., 2010; Madden et al., 2012; Pierpaoli et al., 2001; Song et al., 2003) within the bilateral corpus callosum (portions of the genu, body, and splenium) and right corona radiata, which encompassed portions of the forceps minor, cingulum, and corticospinal tracts and the fronto-occipital fasciculus and anterior thalamic radiations (Fig. 5; Table 5). That is, those individuals who exhibited the most postural sway during the ECF relative to EO condition (i.e., poorer vestibular function) had the lowest ADt within these regions noted above.

**Figure 5:**
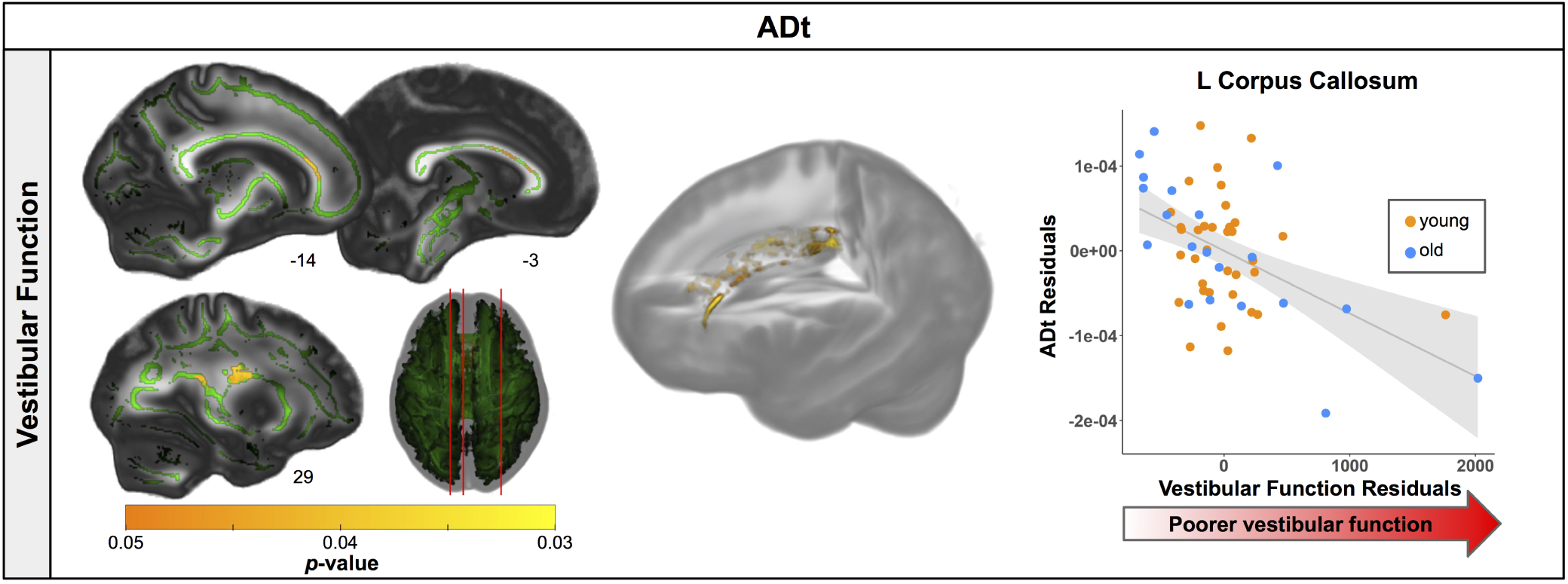
Regions of correlation between ADt and vestibular function scores. Left. Regions showing statistically significant (*p_FWE−corr_* < 0.05) relationships between ADt and vestibular function scores. Warmer colors indicate regions of stronger correlation. Results are shown on the FMRIB58 FA template with the group mean white matter skeleton (green) overlaid. Right. ADt values for the peak result coordinate are plotted against vestibular function score to illustrate an example of the relationship identified by the voxelwise statistical test. The fit line and confidence interval shading are included only to aid visualization of this relationship. We plotted the residuals instead of the raw values here to adjust for the effects of the age and sex covariates included in each model.

**Table 5:**
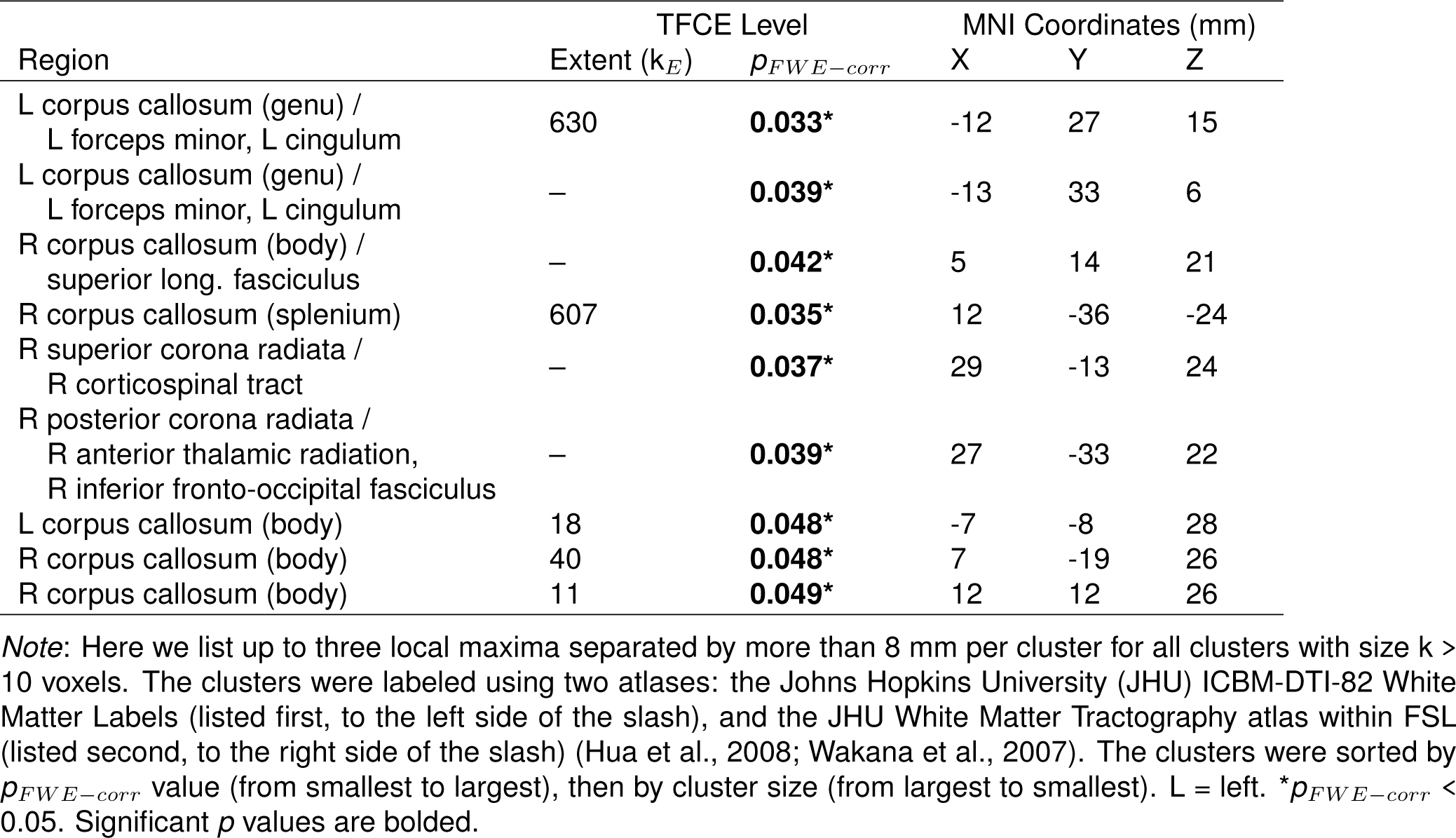
Regions of correlation between ADt and balance scores

| Region | TFCE Level |  | MNI Coordinates (mm) |  |  |
| --- | --- | --- | --- | --- | --- |
| | Extent ($k_E$) | $p_{FWE-corr}$ | X | Y | Z |
| L corpus callosum (genu) /<br>L forceps minor, L cingulum | 630 | **0.033\*** | -12 | 27 | 15 |
| L corpus callosum (genu) /<br>L forceps minor, L cingulum | — | **0.039\*** | -13 | 33 | 6 |
| R corpus callosum (body) /<br>superior long. fasciculus | — | **0.042\*** | 5 | 14 | 21 |
| R corpus callosum (splenium) | 607 | **0.035\*** | 12 | -36 | -24 |
| R superior corona radiata /<br>R corticospinal tract | — | **0.037\*** | 29 | -13 | 24 |
| R posterior corona radiata /<br>R anterior thalamic radiation,<br>R inferior fronto-occipital fasciculus | — | **0.039\*** | 27 | -33 | 22 |
| L corpus callosum (body) | 18 | **0.048\*** | -7 | -8 | 28 |
| R corpus callosum (body) | 40 | **0.048\*** | 7 | -19 | 26 |
| R corpus callosum (body) | 11 | **0.049\*** | 12 | 12 | 26 |

*Note:* Here we list up to three local maxima separated by more than 8 mm per cluster for all clusters with size $k > 10$ voxels. The clusters were labeled using two atlases: the Johns Hopkins University (JHU) ICBM-DTI-82 White Matter Labels (listed first, to the left side of the slash), and the JHU White Matter Tractography atlas within FSL (listed second, to the right side of the slash) (Hua et al., 2008; Wakana et al., 2007). The clusters were sorted by $p_{FWE-corr}$ value (from smallest to largest), then by cluster size (from largest to smallest). L = left. \* $p_{FWE-corr} < 0.05$ . Significant $p$ values are bolded.

### Multiple Regression to Fit the Best Model of Vestibular Function Scores

We used a stepwise multivariate linear regression to compare the predictive strength of the neural correlates of vestibular function score identified above. We entered each participant’s vestibular function score as the outcome variable, and their left supramarginal gyrus cortical thickness, left postcentral gyrus gyrification index, and left corpus callosum ADt, as well as age and sex as predictors. The stepwise regression returned a model containing all of these predictors except for sex. That is, the combination of these brain metrics and age (rather than any given metric on its own) best predicted the vestibular function scores (i.e., produced the model with the smallest AIC; Table 6).

**Table 6:**
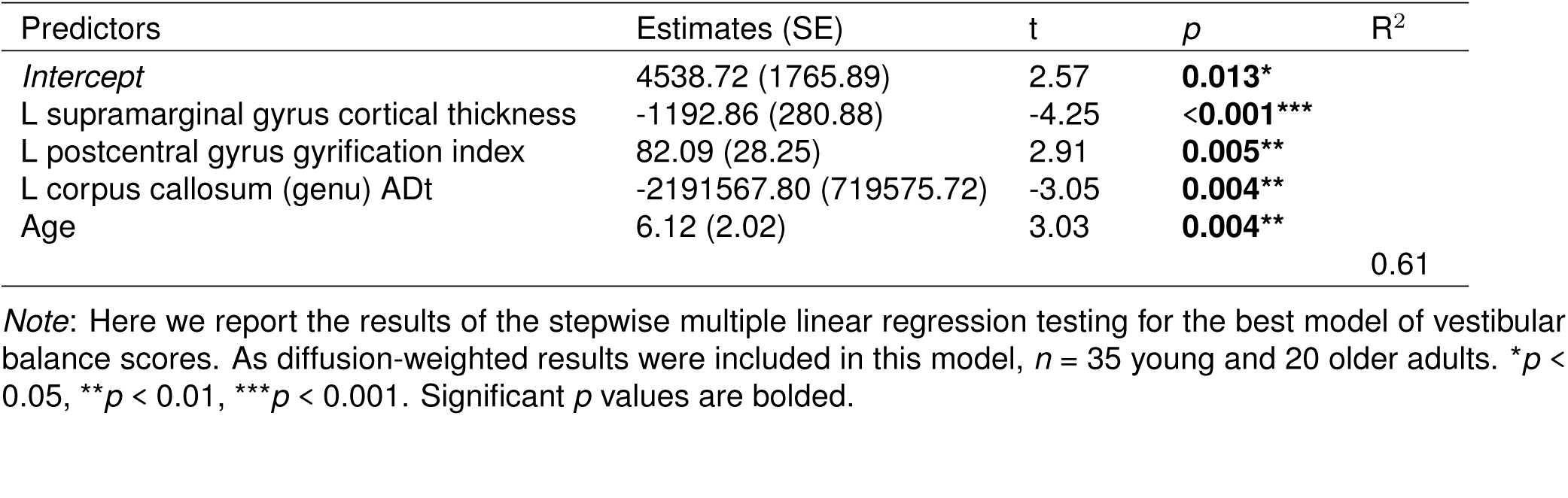
Stepwise multiple regression results for the best model of vestibular balance scores

## Discussion

We identified age group differences for two of the three balance scores, i.e., higher proprioceptive reliance and poorer vestibular function scores for older adults. This indicates that, compared with young adults, older adults rely more heavily on proprioceptive inputs for maintaining balance, and have poorer vestibular function. We also observed multiple significant relationships between brain structure and balance scores. Thinner cortex (i.e., ”worse” brain structure) in regions related to multisensory integration correlated with greater reliance on visual inputs for balance. Higher gyrification index (i.e., more ”youth-like” brain structure) within the sensorimotor and parietal cortices correlated with greater reliance on proprioceptive inputs for balance. Thinner cortex in regions related to vestibular function and lower ADt (i.e., ”worse” brain structure) in the superior-posterior corona radiata and across the corpus callosum were correlated with poorer vestibular function. Higher gyrification index (i.e., more ”youth-like” brain structure) in the sensorimotor, parietal, and frontal cortices was also correlated with poorer vestibular function. These results provide greater understanding of the structural correlates of standing balance control and highlight potential targets for future interventions.

### Age Differences in Balance Scores

Older adults exhibited comparatively more difficulty standing on a foam compared to a firm surface (i.e., higher proprioceptive reliance scores) and during the ECF versus EO condition (i.e., poorer vestibular function scores). There were no age group differences in vision scores. Visual reliance scores (sometimes referred to as a Romberg Quotient) are usually higher for older compared with young adults (e.g., Doyle et al., 2004), though at least one study has reported a lack of age differences in the Romberg Quotient (Lê and Kapoula, 2008). Similar to our results, previous work has identified the greatest postural sway for older compared with young adults when a compliant (e.g., foam) surface is introduced (e.g., Choy et al., 2003; Woollacott et al., 1986). Thus, compared with the young adults, the older adults here may have relied similarly on visual inputs but more so on proprioceptive information for controlling their balance. Though here we interpret higher visual and proprioceptive scores as indicative of greater reliance on these systems for balance, the interpretation of these scores may be more complicated. These scores might index sensory reweighting and integration more so than “reliance” on one sensory system. For example, an increase in postural sway between the EO and EC conditions cannot be attributed only to reliance on visual inputs for balance. It could also indicate difficulty upweighting and properly integrating afferent proprioceptive and vestibular information (Kalron, 2017). Aging has a negative impact on sensory reweighting and integration processes (Colledge et al., 1994; Stelmach et al., 1989; Teasdale et al., 1991; Woollacott et al., 1986). For example, when visual or proprioceptive inputs are removed or altered and then reintroduced, young adults can adapt rapidly and reduce their postural sway, whereas older adults exhibit more postural sway and less adaptation when a new or additional sensory channel is initially added (Hay et al., 1996; Teasdale et al., 1991). Thus, the higher proprioceptive reliance and poorer vestibular function scores we observed for older adults might be due in part to greater difficulty with sensory integration.

### Brain Structure Correlates of Visual Reliance Scores

Across both age groups, thinner cortex within the right cingulate gyrus, precuneus, and lingual gyrus was associated with higher visual reliance scores. Those who exhibited the greatest increase in postural sway between conditions with the eyes closed versus open had the thinnest cortex in these regions. These brain regions do not relate specifically to visual function, but instead play a role in multisensory processing including attentional control, internally-directed cognition, and task engagement (posterior cingulate cortex; Pearson et al., 2011), integration of information and perception of the environment (precuneus; Cavanna and Trimble, 2006), and spatial memory (right lingual gyrus; Sulpizio et al., 2013). It could be that greater reliance on visual inputs is due in part to poorer proprioceptive and vestibular function, and / or brain structure subserving the proprioceptive and vestibular systems (e.g., poorer brain structure in these multisensory processing areas). Thus, individuals may downweight these two systems and rely more on the visual system for balance when all three sensory inputs are available.

This finding could also have been related to sensory integration processes more generally. Poorer brain structure in these multisensory integration regions could have contributed to slower, less effective integration of proprioceptive and vestibular inputs to maintain balance in the absence of visual cues. This would then result in more sway when vision was removed (i.e., higher visual reliance scores). It should be noted that we anticipated better structure (i.e., thicker cortex) in visual processing regions for individuals who typically rely more on vision for balance, due to experience-dependent plasticity processes (May, 2011); however, we did not identify any relationships between canonical visual processing areas and visual reliance scores.

### Brain Structure Correlates of Proprioceptive Reliance Scores

Higher gyrification indices within portions of the left sensorimotor, parietal, supramarginal, paracentral, frontal cortices and precuneus were associated with higher proprioceptive scores (i.e., more difficulty on foam versus firm). Interestingly, the sensorimotor cortex cluster (where the strongest brain-behavior relationship occurred) was located in the cortical region specifically related to lower limb sensorimotor function. Gyrification index generally declines with aging (Cao et al., 2017; Hogstrom et al., 2013; Lamballais et al., 2020; Madan, 2021; Madan and Kensinger, 2018); lower gyrification indices may indicate poorer regional brain structure, i.e., less cortex buried within the sulcal folds (Luders et al., 2006). Thus, it follows that lower gyrification index in a region specifically related to processing lower limb somatosensory information would be associated with less reliance on proprioceptive inputs for balance. As described above, it could be that poorer structure in the brain regions primarily associated with processing one type of sensory information (e.g., proprioceptive) correlates with less reliance on that system and more reliance on other systems (e.g., visual) for maintaining balance.

### Brain Structure Correlates of Vestibular Function Scores

Thinner cortex within two regions encompassing portions of the left supramarginal and postcentral gyri and the bank of the left superior temporal sulcus was associated with poorer vestibular function scores. Stated differently, those individuals who exhibited comparatively more postural sway during the ECF compared to the EO condition also had the thinnest cortex in these regions. These brain regions contribute to vestibular processing and are consistent with vestibular networks identified by our prior functional MRI work (Hupfeld et al., 2020, 2021b; Noohi et al., 2017, 2019) as well as meta-analyses identifying vestibular cortex (Lopez et al., 2012; zu Eulenburg et al., 2012). The supramarginal gyrus is also thought to contribute to proprioception (Ben-Shabat et al., 2015), whole body spatial orientation (Fiori et al., 2015; Kheradmand et al., 2015), and integration of vestibular inputs with visual and proprioceptive information (Ionta et al., 2011). This portion of the temporal sulcus contributes to sensory integration (particularly of audiovisual inputs; Hein and Knight, 2008; Vander Wyk et al., 2009). Thus, it is logical that those with the poorest brain structure (i.e., the thinnest cortex) in these brain regions specifically related to vestibular and multisensory processing also encounter the most difficulty standing with minimal postural sway during a balance condition that specifically tasks the vestibular system.

Higher gyrification indices within parts of the left sensorimotor, parietal, supramarginal, paracentral, frontal cortices and precuneus were associated with poorer vestibular function scores (i.e., more difficulty during ECF compared to EO). This relationship between higher gyrification index and poorer vestibular function is seemingly contradictory. However, as opposed to the relationship described above between thinner vestibular cortex and poorer vestibular function, resulting brain regions for this relationship did not include the vestibular cortices (Lopez et al., 2012; zu Eulenburg et al., 2012). Instead, the strongest relationship between higher gyrification index (i.e., more ”youth-like” brain structure) and poorer vestibular function occurred in the medial pre- and postcentral gyri, which are related to axial and lower limb sensorimotor processing. It could be that those with poorer vestibular function rely more heavily on other brain regions and sensory systems for balance as a compensatory mechanism. However, it should also be noted that the interpretation of gyrification index may be more complex, as a recent study identified relationships between better cognitive function and both higher and lower gyrification index in normal aging and Parkinson’s disease (Chaudhary et al., 2020).

Poorer vestibular function scores were associated with lower ADt within the bilateral corpus callosum and right corona radiata, which encompassed portions of the forceps minor, cingulum, and corticospinal tracts and the fronto-occipital fasciculus and anterior thalamic radiations. Those who exhibited the greatest increases in postural sway between the ECF and EO conditions also had the lowest ADt in these regions. Lower ADt is hypothesized to indicate accumulation of debris or metabolic damage (Madden et al., 2012), axonal injury and subsequent gliosis (Pierpaoli et al., 2001; Song et al., 2003), or disrupted macrostructural organization (Bennett et al., 2010). Across the brain, ADt was largely lower for the older compared with young adults in this dataset (Hupfeld et al., 2021a). Thus, it is logical that lower ADt in these white matter tracts related to interhemispheric communication and motor function would relate to poorer vestibular function.

### Lack of Age Differences in Brain-Behavior Relationships

It is somewhat surprising that we did not identify age differences in the relationship between brain structure and balance. One previous study reported relationships between brain structure and balance for older but not younger adults (Van Impe et al., 2012). In our prior work on this dataset (Hupfeld et al., 2021a), we identified multiple relationships between brain structure and dual task walking for older but not young adults. It is worth noting that this is a group of high functioning older adults in relatively good health, thus, the balance tasks used here may not have been sufficiently biomechanically challenging or cognitively-demanding for age differences in brain-behavior relationships to emerge. If we had incorporated a secondary cognitive task, perhaps we would have found age group differences. Performing a secondary cognitive task has been found to disproportionately affect older adults (e.g., increasing sway variability by 5% for young adults but 37% for older adults; Maylor et al., 2001). An executive function secondary task would have required greater contributions from the prefrontal cortex. Given the large body of literature reporting age-related differences in frontal cortex structure (Fjell et al., 2009; Lemaitre et al., 2012; Salat et al., 2004; Thambisetty et al., 2010), and that balance may require greater attentional control in older age (Dault et al., 2001a,b; Doumas et al., 2009; Huxhold et al., 2006; Rankin et al., 2000), a task with a more challenging cognitive component may have resulted in a correlation between prefrontal cortex structure and balance for the older but not the younger adults.

### Limitations

By using a cross-sectional approach, we could not track concurrent changes in brain structure and balance over time. This approach prevented us from testing whether increased reliance on vision and proprioception over time – in compensation for longitudinal declines in vestibular function – could result in neuroplastic changes in the brain regions responsible for processing these inputs. In addition, the vestibular score did not fully isolate vestibular from proprioceptive contributions; as we compared a foam condition (ECF) to EO, the vestibular score incorporated both proprioceptive and vestibular challenges. Future work could probe additional balance conditions such as a full NeuroCom Sensory Organization Test (SOT), which includes visual conflict conditions. We did not examine other balance outcome variables, such as sway range or velocity. Lastly, in the current acquisition protocol we had a single-shell diffusion sequence. Future studies should consider a multi-shell sequence for a more robust estimation of the free water fraction.

### Conclusions and Future Directions

We identified relationships between regional brain structure (cortical thickness, gyrification index, and ADt) and balance scores indicative of reliance on visual and proprioceptive inputs and vestibular function. Understanding which brain regions contribute to different aspects of balance could be useful in developing future interventions. tDCS, a form of noninvasive brain stimulation, has been demonstrated to augment balance performance and training for both young and older adults (Hupfeld et al., 2017a,b; Kaminski et al., 2016; Yosephi et al., 2018). Uncovering how brain structure relates to balance function could help identify regions to target with tDCS. This is a promising future intervention, with some evidence showing that tDCS affects brain function (Pupíıková et al., 2021) and neurochemicals (Heimrath et al., 2020), and produces effects that may last for months post-stimulation (Vestito et al., 2014).

## Author Contributions

KH led the initial study design, collected and preprocessed all of the neuroimaging and gait data, conducted all statistical analyses, created the figures and tables, and wrote the first draft of the manuscript. OP and HR consulted on DWI preprocessing and contributed to manuscript preparation. CH consulted on the design and analysis of the gait assessments. RS oversaw project design and led the interpretation and discussion of the results. All authors participated in revision of the manuscript.

## Acknowledgments

The authors wish to thank Aakash Anandjiwala, Justin Geraghty, Pilar Alvarez Jerez, and Alexis Jennings-Coulibaly for their assistance in subject recruitment and data collection, as well as Sutton Richmond for his help in applying the signal drift correction to the diffusion-weighted data. The authors also wish to thank all of the participants who volunteered their time, as well as the McKnight Brain Institute MRI technologists, without whom this project would not have been possible.

